# Intrinsic neural activity predisposes susceptibility to a body illusion

**DOI:** 10.1101/2021.09.18.460883

**Authors:** Tzu-Yu Hsu, Ji-Fan Zhou, Georg Northoff, Su-Ling Yeh, Timothy Joseph Lane

## Abstract

Susceptibility to the rubber hand illusion (RHI) varies. Thus far, however, there is no consensus as regards how to explain this variation. Previous studies, focused on the role of multisensory integration, have searched for neural correlates of the illusion. Those studies, however, have failed to identify a sufficient set of functionally specific neural correlates. An alternative explanation of the illusion is that it results from demand characteristics, chiefly variability in the disposition to respond to imaginative suggestion: the degree to which intrinsic neural activity allows for a blurring of boundaries between self and external objects. Some evidence suggests that frontal α power is one means of tracking neural instantiations of self; therefore, we hypothesized that the higher the frontal α power during eyes-closed resting state, the more stable the self. As a corollary, we infer that the more stable the self, the less susceptible are participants to a blurring of boundaries—to feeling that the rubber hand belongs to them. Indeed, we found that frontal α amplitude oscillations negatively correlate with susceptibility. Moreover, since α and δ oscillations seem to be associated in pathological states that allow for a blurring of boundaries between self and external objects, we conjectured that the high frontal α power observed in low-RHI participants is modulated by δ frequency oscillations. Indeed, we found this to be the case. Based on our findings we propose that the two explanatory frameworks might be complementary: that is, the neural correlates of multisensory integration might be necessary for the RHI, but a sufficient explanation requires investigation of variable intrinsic neural activity that acts to modulate how the brain responds to incompatible sensory stimuli.

**Highlights:**

~Intrinsic frontal α power negatively correlates with susceptibility to the RHI.
~Intrinsic α power modulated by δ oscillations varies with susceptibility to the RHI.
~Sufficient explanation of RHI requires understanding of intrinsic neural dispositions that regulate the boundary between self and the external world.

## Introduction

The body is malleable, albeit more for some than for others. Even lifeless objects can be experienced as belonging to self. Tastevin (1937) suggested that ersatz hands can be experienced as belonging to self when participants simply look at them. Then under experimental conditions, Botvinick and Cohen (1998) formalized and replicated induction of this “rubber hand” illusion (RHI). Subsequently, the RHI has been extensively replicated, by various techniques, multiple times, and for many purposes (Braun 2018; de Vignemont 2018; Ehrsson 2009a 2012; Makin et al. 2019; Holmes & Ehrsson 2008; Riemer 2019; Tsakiris 2010, 2011, 2012). But we do not yet know what intrinsic neural activity makes possible this illusion.

On the standard version of the RHI, an artificial hand is experienced as belonging to self when subjects see it being stroked, concurrent with feeling strokes applied to the occluded, real hand. But when visual and tactile sensations are asynchronous—the common control condition—the illusion either fails to occur or is less vivid (Tsakiris 2010). The RHI paradigm has become a cornerstone of investigations into the science of self, and the distinctive ownership illusion tends to be explained as the result of multisensory integration of a type whereby vision prevails over other senses (Ehrsson 2012, Tsakiris 2017, Braun et al. 2018).

Subjective report evidence is adduced from questionnaires that generate data amenable to psychometric analyses (e.g., Longo et al. 2008, Romano et al. 2021). Behavioral or physiological evidence is derived from measures like proprioceptive drift, skin conductance, time of onset or duration, and temperature (e.g., Critchley et al. 2021, de Haan 2017, Ehrsson et al. 2004, Lane et al. 2017, Yeh et al. 2017). The multisensory integration hypothesis suggests that neural activity that mediates the experience seems to involve the premotor cortex (PMC), the intraparietal sulcus (IPS), the anterior insula, and the sensorimotor cortex (Bekrater-Bodmann et al. 2014; della Gatta et al. 2016; Ehrsson et al. 2004; Kammers et al. 2009; Limanowski et al. 2014; Lira et al. 2018; Petkova et al. 2011; Peviani et al. 2018). It has not yet been possible, however, to identify a sufficient set of functionally specific neural correlates; in fact, even when employing the higher time resolution of EEG, results have been inconsistent (for a summary of the relevant studies see Rao and Kayser 2017).

In addition, several confounds have contributed to the failure to find neural correlates of consciousness (NCCs) for the RHI (Swinkels et al. 2020). Arguably the most disconcerting, purported confound implicates demand characteristics, the ability of subjects to grasp and comply with intentions suggested by experimental design. As this applies to the RHI, it might be that multisensory integration is necessary but not sufficient to explain the illusion; subjects with a certain trait seem better able than others to confabulate the ownership subjective experience and report (Lush et al. 2020, 2021; Dienes et al. 2020; Marotta et al. 2016). That is to say, although investigations of the RHI NCCs have some explanatory value, a sufficient explanation will require inclusion of neural predispositions.

On the psychological level, the proposal is that we should attend more to what some researchers refer to as “trait phenomenological control” (Lush et al. 2020)—or, prosaically, suggestibility. The point is not that the experiences are any less real; the point is that the relevant NCCs likely omit certain mechanisms, akin to those that underlie suggestibility. One implication of this idea is that resolution of several explanatory problems, such as the variability of RHI or other body illusions (Botvinick 2004, Durgin et al. 2007, Ehrsson et al. 2004, Ehrsson 2012, Kanayama et al. 2009, Lloyd 2007, Mariotta et al. 2016, Yeh et al. 2017), will likely involve a model that includes suggestibility.

In point of fact performance on a hypnotizability scale (Lush et al. 2018) predicts much of the observed variability (Lush et al. 2020). This finding implies that suggestions implicit in the paradigm might be necessary to precipitate not only the illusory experience, but also the corresponding physiological responses and brain activity. Our purpose here is to take this concern with suggestibility (or hypnotizability) seriously and seek to determine whether there might be neural evidence of a trait that predisposes subjects to experience the RHI.

Consider that hypnotizability is a normally distributed, stable trait, but for those who exhibit this trait, response to suggestion does not require hypnotic procedures (Lush et al. 2020). What matters is the disposition to respond to imaginative suggestion, and in this instance the imaginative suggestion is that the rubber hand belongs to you. But what does responsiveness to imaginative suggestion mean? A clue is found in the fact that hypnotizability’s most reliable correlate is “absorption,” which can be understood as the disposition to undergo experiences for which *boundaries between self and object are blurred* (Kihlstrom 2016). Based upon prior findings concerning this correlate, we conjecture that one mechanism relevant to the RHI that helps to explain variability, but that has heretofore remained unexplored, involves neural instantiation of self.

Fortunately, it has become possible to get a handle on neural instantiations of self by attending to the brain’s intrinsic activity (Gusnard et al. 2001, Smallwood and Schooler 2015, Qin et al. 2016). In other words, in order to explain the RHI, we propose that the explanatory framework must be expanded beyond NCCs and include neural predispositions. Indeed, among the more prominent types of neural activity that help track instantiation of self is resting state α (alpha) power, RSAP (Kraus et al. 2021). Indeed, resting state—the brain’s spontaneous, intrinsic activity—frontal α power can be used to predict whether subjects will identify external stimuli as self-related (Bai et al. 2016, Lane et al. 2016).

Clearly intrinsic α power can be employed to track numerous phenomena (e.g., Clayton et al. 2018, Hanslmayr et al. 2007, Hutchinson 2021, Piantoni et al. 2017, Romei et al. 2008 a, b, Samaha and Postle 2015, Thut et al. 2006), so our claim is not that it represents or is the neural realization of self. But it can serve as one proxy whereby self-related neural activity can be tracked (Lane 2020). Motivated by these findings, and in order to probe more deeply into data that suggest hypnotizability helps explain the RHI variability, we analyzed EEG data, focusing on variability in whether participants feel the ersatz hand *belongs* to self (Lane 2015). Our *hypothesis* is that frontal RSAP can be used to track the degree to which a subject’s boundaries between self and an inanimate object (viz., the rubber hand) are susceptible to blurring. To cast this idea in colloquial terms: if we take frontal RSAP as a means of tracking the persistence of self-boundaries, despite the introduction of multisensory stimuli designed to trick the brain’s integrative tendencies, the greater the RSAP, the less likely a brain is to succumb to the illusion. In short, the greater the RSAP, the less likely a participant is to feel the rubber hand belongs to self.

What is more, lower frequencies often modulate higher frequencies (Brookes et al. 2011, Buzsáki & Draguhn 2004, Conolty et al. 2006), enabling integration of information from distal brain regions at low energy costs (Buzsáki 2019, Freeman and Rogers 2002; Vanhatalo et al. 2004). Based on the assumptions that multisensory integration continues to play some role in the RHI, and that information integration across distributed regions is necessary for the realization of conscious experiences—for which illusions are paradigmatic—we further conjectured that RSAP is modulated by the phase of lower frequency oscillations. When considering where to look among lower frequencies, we examined other states wherein boundaries between self and object are blurred, in particular schizophrenia (Lane 2014). Indeed, it appears to be the case that participants who are prone to psychosis are more likely than are controls to have vivid RHI experiences (Peled et al. 2000, Thakkar et al. 2011, Germine et al. 2013). What is more, during eyes closed (EC) resting state, schizophrenia patients evince relative increases in lower frequency power, along with decreases in the absolute α band power (Newson and Thiagarajan 2018, Ranlund et al. 2014). And since the ratio of δ (delta) to α—increased δ and decreased α—is associated with psychosis, we focused on δ (Howells et al. 2018).

In sum, we investigated EEG data, examining both RSAP and phase-amplitude coupling for δ and α frequency oscillations, focusing on the EC resting state, both prior to and after the RHI task. We *hypothesized* that time frequency analysis would show these two frequency bands to be related to RHI susceptibility. In short, we analyzed EC resting state data, while giving special attention to δ and ± along with questionnaires designed to assess the degree to which participants report feeling that the ersatz hand belongs to them.

## Methods

### Participants

Twenty-four right-handed college students from the National Taiwan University community participated in this study. All had normal or corrected-to-normal vision and all were neurologically unimpaired. This study was approved by the Research Ethics Committee, National Taiwan University. All participants gave informed consent.

### Experimental Procedures

This study comprised two stages: In Stage One the standard RHI induction task is employed (see below). For this stage participants indicated illusion onset by pressing their feet to pedals, in a manner that we used in our prior investigations (Lane et al. 2017, Yeh et al. 2017). In Stage Two, EEG experimental procedures are employed (see below). The purpose of dividing the experiment into two stages was to first identify distinct subsets of participants—high and low susceptibility. Marking this distinction was in preparation for the second stage, in which the main hypothesis—viz. high and low would evince intrinsic neural activity differences—would be tested.

### Stage One: The standard RHI induction task

Participants were tested, individually, in a small, quiet room. The experimenter sat in front of the participant, who was seated with both hands placed on a table top. Participants were asked to insert their hands into a black cardboard tube, so that they would be hidden from view. A towel was placed over the tube and in such a way that it would conceal participants’ elbows and forearms, as well as the space separating the rubber hand from the body (see Lane et al. 2017, Fig 1). Then, the experimenter proceeded to use two paintbrushes to stroke corresponding fingers of rubber and biological hands, at an approximate rate of one per two seconds. Because evidence suggests right hemispheric dominance for the experience of body ownership (Ocklenburg, et al., 2011), strokes were applied to the real left hand and a corresponding left, rubber hand. The procedure ceased either at 15 s after illusion onset or after three minutes, for those participants who did not experience the illusion. It was in this way that we identified the distinct subsets of susceptibility, in preparation for stage two.

**Figure 1.**
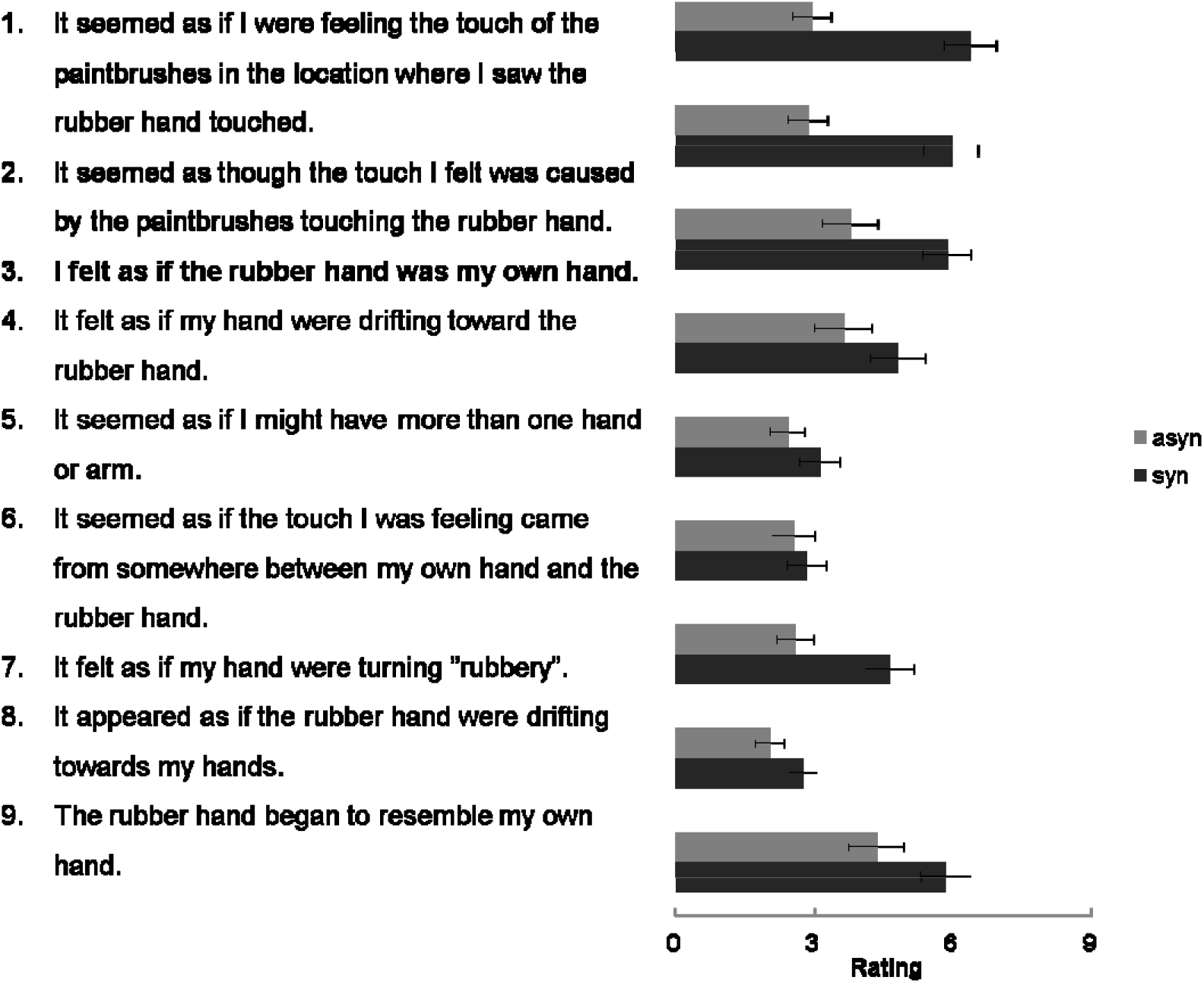
The questionnaire used in this study (adopted from Botvinick and Cohen, 1998). Item #3 concerns ownership for the rubber hand. The right panel is the mean scores for each item.

Two conditions were administered for each participant: synchronous and asynchronous stroking, with the order counterbalanced across participants, and the asynchronous condition was treated as the control. In both conditions the experimenter stroked the rubber hand and participants’ real hands, attempting to induce the RHI. Participants were required to keep looking at the rubber hand and avoid postural adjustments. After stroking of the hands was completed, participants were asked to fill out questionnaires.

The main questionnaire used for this experiment was adapted from Botvinick and Cohen (1998). It was used to evaluate whether, or the degree to which, the illusion was experienced in the synchronous condition, relative to the asynchronous. Responses for each item were indicated on a nine-point scale, from 1 to 9. Questionnaire scores for each participant were assessed by subtracting synchronous from asynchronous scores. Because in this stage we used the standard control condition for assessing susceptibility, it was this questionnaire data that we matched to the EEG data collected in stage two.

### Stage Two: EEG Experimental procedure

All participants underwent the RHI task procedure; each time the stroking continued for one minute. The duration of stroking was one minute, both because of our findings from Stage One, and because other investigations of the RHI have found that it can occur in less than 12 seconds (Ehrsson, Spence, & Passingham, 2004; Arzy, Overney, Landis, & Blanke, 2006; Tsakiris & Haggard, 2005; Tsakiris et al., 2007). Moreover, because the standard RHI induction task in Stage One enabled us to distinguish between two groups—those who are susceptible and those who are not—we did not use the usual control condition: that is, only synchronous stroking was employed, five times for each participant.

As for EEG measurements of intrinsic neural activity, a total of one minute of eyes-open (EO) and one minute of eyes-closed (EC) resting activity were measured, both before and after the induction of RHI. Both EO and EC were sustained for 20 s, and repeated three times: in each instance we began with EC and then alternated between EC and EO (viz. EC, EO, EC, EO, EC, EO) three times. During the EC, participants were instructed to close their eyes and relax without thinking of anything in particular; whereas, during EO, participants were required to fixate on a cross. Importantly, one minute of recording is sufficient to detect the low frequency, intrinsic oscillations, that are the target of our investigation (Lu et al. 2007; Karamacoska, Barry, & Steiner 2017; Mantini et al. 2007).

### EEG recording and Analysis

EEG activity was recorded with Ag/AgCl electrodes mounted in an elastic cap (Electrocap International) using a 64-electrode arrangement, following the International 10-20 System. Two additional electrodes were referenced to the left and right mastoid. Vertical eye movements were recorded from electrodes above and below the right eye and horizontal electrooculograms were also recorded from electrodes at the outer canthi. Electrode impedances were kept below 10 kΩ for all electrodes, and amplifier bandpass was 0.1-100 Hz. Data were recorded with Neuroscan 4.2 software, with a sampling rate of 1000 Hz.

All data analysis was performed off-line using EEGLAB (Delorme & Makeig, 2004) and custom MATLAB (MathWorks) scripts. The continuous EEG was then segmented into 20-second-long epochs. ICA was applied to remove eye-movement induced artifacts (blinks or saccades). Also, epochs containing excessive noise or drift (± 100 μV) at any electrode were excluded. The power of the signals from each channel was computed in 1 and 40 Hz (Roach and Mathalon, 2008) using Morlet wavelet convolution with 3 cycles and 1 Hz step. Finally, power was calculated by converting the signal to a decibel (dB) scale, by multiplying logarithm (10*log 10[power]). All contrast between conditions was tested with nonparametric permutation testing; this corresponds to a cluster-level threshold of p < 0.01.

In order to explore cross-frequency coupling during intrinsic activity, i.e., phase-amplitude coupling (PAC) between the pair of frequencies (Canolty et al. 2006), we adopted a modified version of the modulation index (PACz, Cohen, 2014). This was done in order to control for potential confounds and render the data amenable to statistical evaluation (Cohen, 2014). First, the raw signal was separated into α power and δ phase through zero-phase band-pass filtered from 8-13 Hz and 1-3 Hz, respectively; the Morlet wavelet was then applied to estimate the amplitude of α frequency oscillation and the phase of δ range oscillation. Second, the modulation index (Canolty et al. 2006) was used to identify coupling between α amplitude and δ phase. The analysis was done with selected electrodes from anterior sites (AF3, AF4, F1, Fz, and F2); anterior sensors were selected in order to test our hypothesis concerning frontal α activity. Third, surrogate signals (n=1000) were computed to create randomly permuted power time series distribution of PAC values. We did not use the raw PAC values; instead, PACz was estimated through PAC value that was normalized with respect to randomly permuted PAC values. The normalized PAC—the PACz—was normalized by z values (Cohen, 2014). Therefore, we adopt the z-score to identify PACz’s α threshold of p <0.05. To control for Type I errors when conducting multiple comparisons, the false discovery rate (FDR) method (Benjamini and Hochberg, 1995; Groppe et al., 2011) was applied—a p-value of less than 0.05 survived FDR-controlling procedures.

## Results

### Stage One: Synchrony/asynchrony manipulations for standard RHI induction

To ensure that we have replicated previous RHI studies, two-way repeated-measures ANOVA with Greenhouse-Geisser adjustment was applied to compare questionnaire scores across synchrony and asynchrony conditions, for each item. For the questionnaire scores from Botvinick and Cohen (1998), a significant main effect of synchrony was found, *F*(1,2,3)=35.547, *p*<.001 (mean scores: synchrony=4.685; asynchrony=3.019) and the main effect of item, *F*(3.487, 80.208)=12.16, *p*<.001. Scores for the synchrony condition were higher than for the asynchrony condition (Figure 1). In addition, a significant interaction between synchrony and item also was found, *F*(3.475, 79.914)=8.143, *p*<.001. Post hoc analysis showed that the scores for most items in the synchronous condition were significantly higher than scores for the asynchronous condition (*p*<.01), except for items four and five (*p*s>.05). Hence, our findings were in line with previous RHI studies.

Of most direct relevance to our investigation, scores for the first three items were in line with findings from the original investigation, including item three, the question that probes the sense of belonging or ownership (Botvinick & Cohen 1998). For those who are susceptible to the RHI, significant synchrony-induced RHI ownership values were evinced. We adopted subtracted scores (synchronous - asynchronous) to represent questionnaire results for the EEG analysis that follows.

### Stage Two: The relationship between intrinsic neural activity and RHI susceptibility

Intrinsic neural activity for the two conditions was transformed into time-frequency power. We then compared time-frequency power of intrinsic neural activity in EO and in EC conditions, both before and after RHI induction. When comparing pre- and post-RHI induction, no significant difference for intrinsic neural activity was discerned. Accordingly, intrinsic neural activity from before and from after RHI induction were pooled for subsequent analysis, while distinguishing EO from EC conditions.

Our findings for time-frequency power across different electrodes as a function of 1-40 Hz are shown in Figure 2. The magnitude of α frequency power was significantly higher for EC, relative to EO conditions, across whole electrodes. Moreover, and precisely as is to be expected, the right panel in Figure 2 (permutation test, *p*<.01) shows that the magnitude of α frequency power on posterior electrode sites significantly increased during EC.

**Figure 2.**
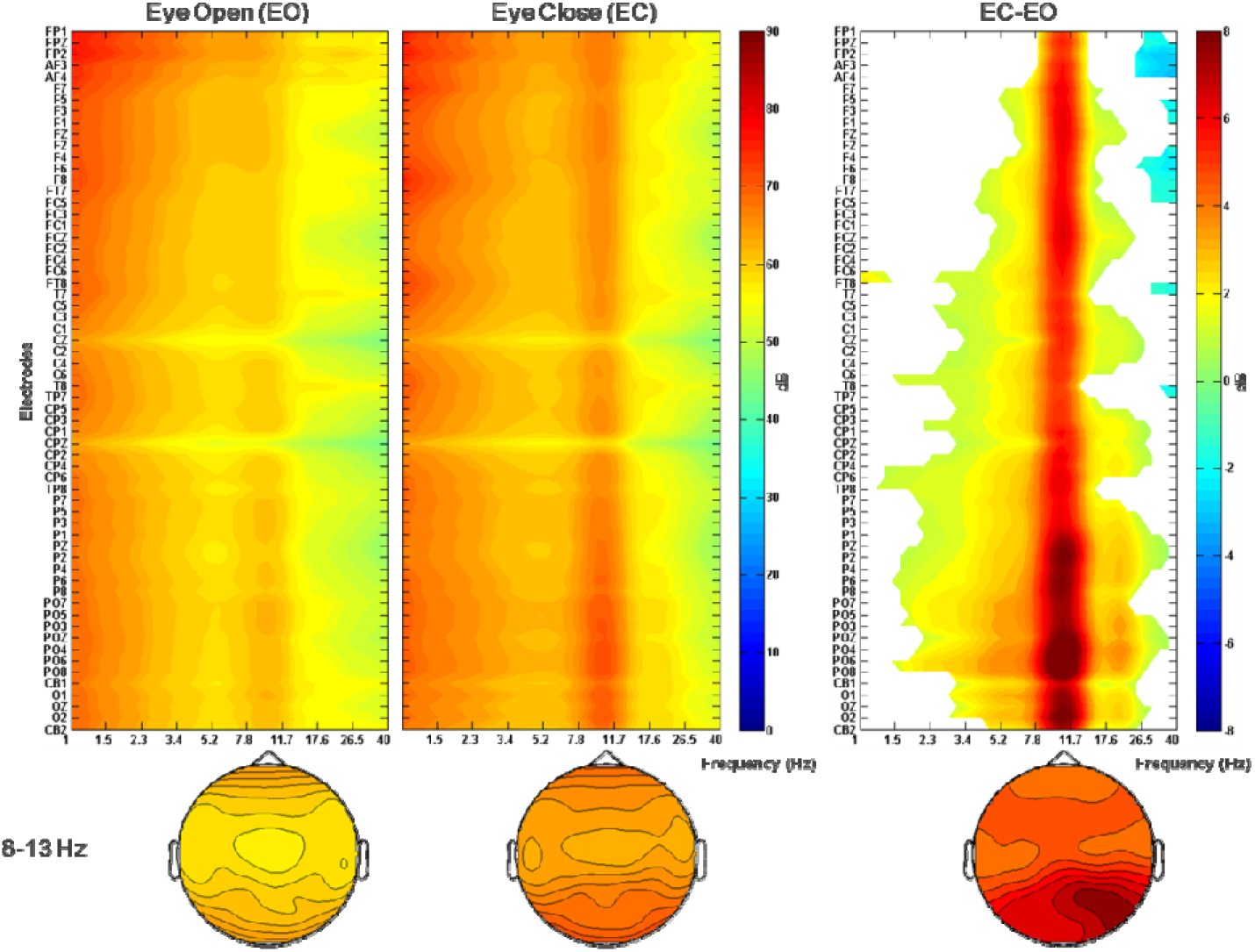
Ground averaged power spectrums during resting states. Power spectrum was ploted as a function of electrodes and is shown for eyes open (EO), eyes closed (EC), and for the difference between EC and EO, from left to right panels. Bottom panels show topography of α frequency power for each condition. α frequency power differences increased in strength, from the frontal to posterior electrodes.

We further examined whether the subjective experience of RHI—in particular the ownership values for item #3—correlates with the amplitude of α frequency power. Accordingly, we conducted a spearman’s rho correlation on the difference scores of ownership in item #3 and the amplitude of α frequency power across central-to-frontal electrodes. The results showed significant negative correlations between ownership scores and amplitude of α power in frontal to central electrodes in the EC condition (*p*<.05, FDR, top panel in Figure 3), but not in the EO condition. To visualize the relationship between the α frequency power and ownership scores in the EC condition, α frequency power for frontal electrodes (i.e. AF3, AF4, F1, Fz, and F2) were averaged, revealing a correlation with ownership scores, *rho*(23)=-.58, *p*<.001. The results showed that α frequency power decreased as ownership scores increased in the EC condition (below, Figure 3).

**Figure 3.**
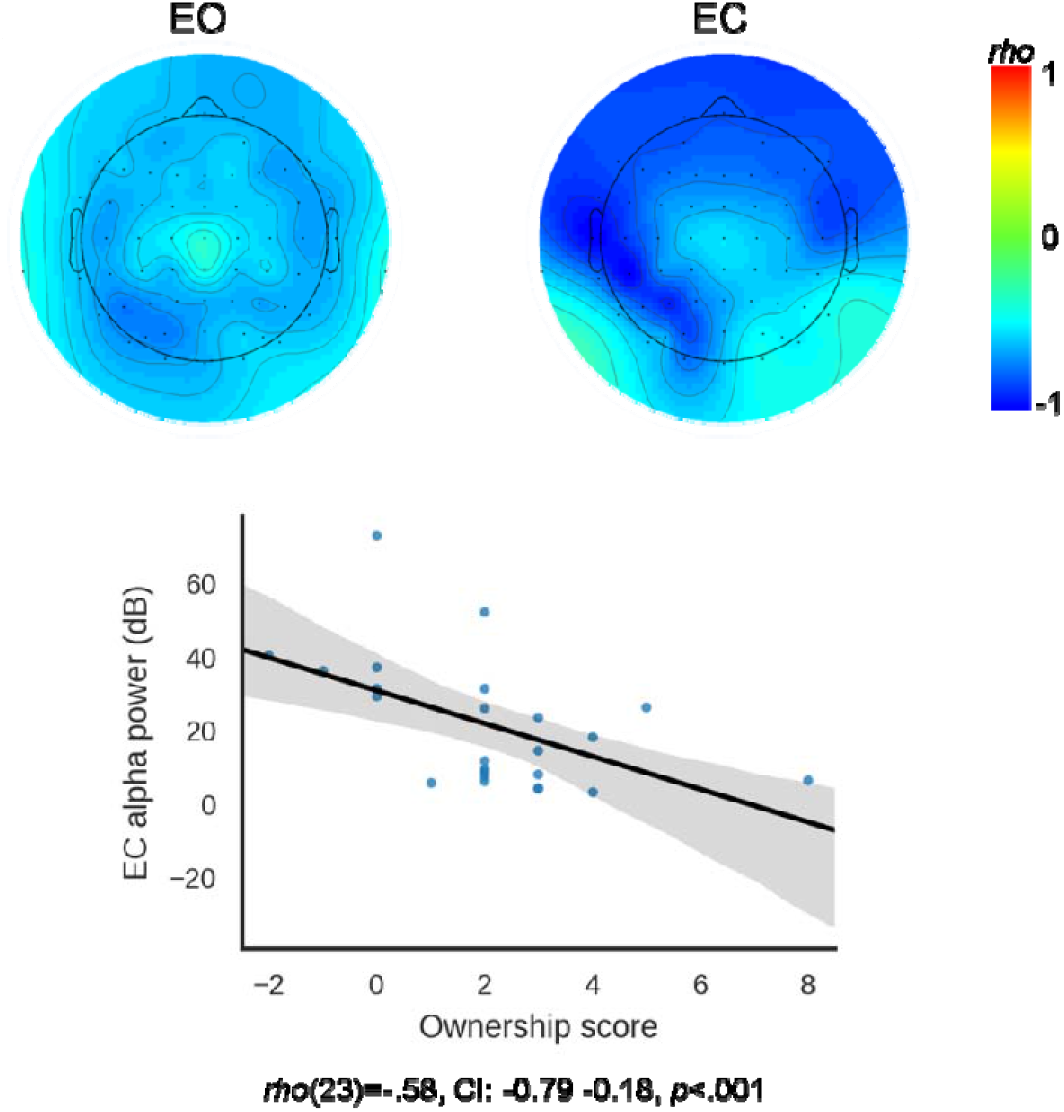
Correlation between ownership scores with alpha power. The top panel showed rho values between ownership scores across electrodes and EO/EC conditions. Significant negative correlation between ownership scores with α power in frontal and central electrode sites was found in the EC condition only. To visualize the negative correlation between the α power and ownership scores, we further combined α power from frontal electrodes (i.e., AF3, AF4, F1, Fz, and F2) in EC condition and did a correlation with ownership scores as shown in the bottom panel.

### Different RHI cohorts evinced different levels of low and high frequency oscillation coupling in the EC condition

As mentioned in the introduction, results of psychosis/psychosis-prone studies in intrinsic neural activities and RHI suggest the possibility of a link between δ and α frequency oscillations, and the RHI. Therefore, we conducted an exploratory analysis by calculating δ phase and α amplitude coupling (i.e., PACz) in the EC condition, and categorized participants into high- and low-RHI groups, based on their ownership scores from item #3 and median split into half. If cross frequency coupling for intrinsic neural activity was irrelevant to RHI susceptibility, there would be no significant PACz relationship between either high-RHI or low-RHI. On the other hand, if PACz either in high-RHI or low-RHI participants is significant, it suggests that underlying, intrinsic neural activity in high-RHI or low-RHI groups differ. Results showed that the PACz for the low-RHI group was significantly higher than zero (PACz=1.986, *p*<.05), whereas PACz for the high-RHI group was not (PACz=0.923, Figure 4). These results suggest the possibility that for the EC condition and in the low-RHI group, communication across δ and α frequency oscillations is heightened.

**Figure 4.**
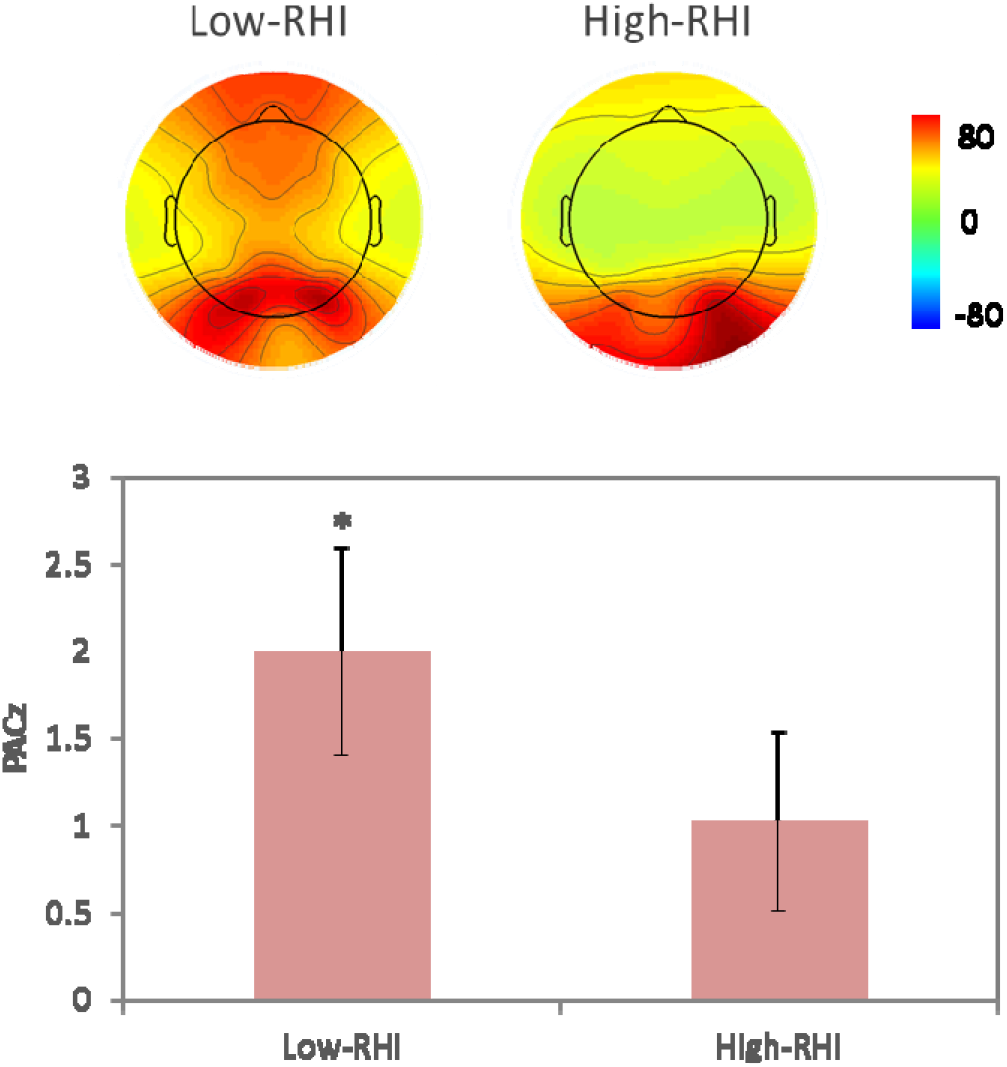
δ phase α amplitude coupling for low- and high-RHI performers in EC condition. Top panel was topography of α frequency power EC condition. Left topography indicates low-RHI and right topography indicates high-RHI scores performers. Averaged PACz in low-RHI performers was significantly larger than Z-score 1.96 (bottom left) whereas averaged PACz in high-RHI performers was not.

## Discussion

To the best of our knowledge, this is the first study to describe an association between the brain’s intrinsic activity and RHI susceptibility. Our findings suggest that temporal, neural predispositions—EC RSAP and phase amplitude coupling for δ and α frequency oscillations—might constrain susceptibility to the RHI. Our findings not only help explain why multi-sensory integration does not affect everyone in the same way, it also lends support to the conjecture that “trait phenomenological control”, or suggestibility, can help explain the RHI (Lush et al. 2020), and perhaps other body illusions. The neural facility for integrating mismatched sensory stimuli—visual, tactile, and proprioceptive—varies from person to person, and suggestibility, possibly underlain by neural predispositions, helps explain why.

The magnitude of frontal α frequency power oscillations in the EC condition is negatively correlated with the score of item #3 on the RHI questionnaire, “I felt as if the rubber hand was my own hand.” It seems to be the case that frontal, α power might carry information relevant to how malleable the subjective sense of self-own-body is, at least if RHI-like incongruous manipulations are employed. The cross-frequency coupling analysis in the EC condition provides additional data concerning neural predispositions that might be relevant to the conscious experience of one’s own body, by showing that underlying cross frequency communications differ between low- and high-RHI subjects. The high frontal α power observed in low-RHI participants appears to be modulated by δ frequency oscillations. This finding might imply that low-RHI participants are better able to preserve their original subjective sense of their bodies, because that sense is bolstered by communication among distal brain regions.

Other investigators of related phenomena have called attention to the significant role played by the α band power, especially when it involves regions that contribute to self-related processing, like the medial prefrontal cortex, mPFC (Lenggenhager et al 2011). Indeed, several studies have suggested that self-related processing involves intrinsic frontal activity (Ben-Simon et al. 2008, Fingelkurts et al. 2012, Knyazev et al. 2012, Knyazev 2013, Lane 2020, Lehmann et al. 2001). What we are proposing is that to the degree that self-related neural processing tracks the self, this neural processing might be able to serve as a proxy for assessing stability of the subjective sense of one’s own body. That is to say, the more stable the self, the less malleable it is, even in the face of mismatched, synchronous stimuli.

We should add that a simpler hypothesis suggests itself: According to the inhibition-timing hypothesis (Klimesch et al. 2007), α rhythms can play an inhibitory role that includes modulation of the temporal windows whereby divergent stimuli can be integrated into unitary percepts (Bastiaansen et al. 2020). In this respect, in order to explain how multisensory integration is constrained (Cecere et al. 2015), there might be no need to invoke the concept of “self.” Instead, citing the inhibition-timing function of intrinsic α oscillations might be sufficient. But to explain the RHI, we choose the abstract concept, “self,” because α oscillations are likely only one proxy for tracking neural instantiations of the degree to which the boundaries between self and external objects, like rubber hands, can be blurred. Another way to say this is that the embodied self is multiply realizable, so the explanatory concept must be pitched at a level of abstraction above the neural.

In a previous study we found that participants who evince less switch cost and higher attention-shift scores had faster RHI onset times; moreover, those who evince higher attention-shift scores experienced the RHI more vividly (Yeh et al. 2017). The findings suggest that a disposition for performing these executive functions contributes to illusion susceptibility. It follows then that if task switch and attention-shift are subserved by frontal neural activity, it might be that frontal α power plays an inhibitory role, consistent with the inhibition-timing hypothesis. In other words, frontal α power might be inhibiting attention shift and task switch, thereby inhibiting illusion susceptibility. The explanation that we have proposed here, on the other hand, is that EC RSAP and phase amplitude coupling for δ and α frequency oscillations makes the embodied-self more stable, thereby inhibiting illusion susceptibility Further investigations of how intrinsic neural activity facilitates or inhibits susceptibility to body illusions will be required in order to determine which explanatory framework more nearly approximates the truth.

Our intent is not to deny that NCCs for multisensory integration are relevant to the RHI. Instead, we suggest that to treat this illusion as a unitary phenomenon is misleading. Given the coarse standards for assessing whether a participant experiences the illusion, it could well be the case that some instances wherein a person reports vivid experience of the illusion result more from suggestion, while others, more from multi-sensory integration. Subjective reports tend to conflate distinct phenomena: This is to say that how one makes a report in response to the questionnaire should not be assumed to be a report about a singular experience. What we propose is a conditional: if participants are making reports about a singular type of experience, multi-sensory integration is not a sufficient explanation. A necessary precondition involves durability of the boundaries between self and the external, and one possible means of tracking this durability involves frontal α power nested in δ oscillations.

We further note that, in broad outline, our findings are consistent with EEG investigations of hypnosis. Previous research has reported that slow wave activity facilitates response to hypnotic suggestion (Jensen et al. 2016). But in fine detail, our findings differ. Unlike those studies we did not find that θ plays a prominent role. Among the possible reasons for this difference, EEG investigations of hypnosis have not been primarily focused on intrinsic activity, and our focus has been on what neural activity might impede—rather than facilitate—the sort of multisensory integration that makes possible the RHI.

Two distinct frameworks have been proposed to explain the several facets of the RHI, including its intersubjective variability. One emphasizes multisensory integration; the other, suggestibility. Although we allow for the possibility that RHI is a heterogeneous phenomenon, we submit that most instances of how this illusion is experienced require both an understanding of the NCCs of multi-sensory integration and an understanding of the intrinsic, neural predispositions that regulate the boundaries between self and the external world. Not all bodies are equally malleable—our findings suggest that frontal α amplitude modulated by δ oscillations contributes to the stability of our sense of where our hands are. If future studies apply this approach to full-body illusions, this intrinsic neural activity might also help to explain our sense of where we are.

## Acknowledgments

Research reported here is supported by funding from Taiwan Ministries of Health and Welfare, Education, and Science and Technology: 110IIT-03, 109TMU-SHH-23, DP2-110-21121-01-N-07-01, MOHW109-TDU-B-212-114007, MOHW110-TDU-B-212-124007, MOST109-2410-H-038-009, 108-2410-H-038-001, 107-2410-H-002-129-MY3. We thank Cheng-Yun Teng and Ting-Yi Lin, for their help in collecting part of the data.

## Author contributions

JZ, TL and SY designed the study; TL, SY and GN contributed to concept development; JZ collected the data; TH and JZ analyzed the data; TH and TL wrote the manuscript; all authors revised and approved the manuscript.

## Data accessibility

Participants did not give consent to their data being shared openly and so the data included in this analysis are only available from the authors upon reasonable request.

## References

Arzy, S., Overney, L. S., Landis, T., & Blanke, O. (2006). Neural Mechanisms of Embodiment. Archives of Neurology, 63(7), 1022. DOI:10.1001/archneur.63.7.1022

Bai, Y., Nakao, T., Qin, P., Chaves, P., Heinzel, A. H., Duncan, N., Lane, T. J., Yen, N-S., Tsai, S-H., Northoff, G. 2016. Resting state glutamate pre-stimulus alpha increase during self-relatedness—A combined EEG-MRS study on rest-self overlap. Social Neuroscience 11, 3, 249–263.

Bastiaansen, M., Berberyan, H., Stekelenburg, J. J., Schoffelen, J. M., Vroomen, J. 2020. Are alpha oscillations instrumental in multisensory synchrony perception? Brain Research 1734, https://doi.org/10.1016/j.brainres.2020.146744

Bekrater-Bodmann R, Foell J, Diers M, Kamping S, Rance M, Kirsch P, et al. (2014) The Importance of Synchrony and Temporal Order of Visual and Tactile Input for Illusory Limb Ownership Experiences - An fMRI Study Applying Virtual Reality. PLoS ONE 9, 1, e87013. https://doi.org/10.1371/journal.pone.0087013

Benjamini, Y., & Hochberg, Y. (1995). Controlling the False Discovery Rate: A Practical and Powerful Approach to Multiple Testing on JSTOR. Journal of the Royal Statistical Society. Series B (Methodological), 57(1), 289–300.

Ben-Simon, E., Podlipsky, I., Arieli, A., Zhdanov, A., & Hendler, T. (2008). Never Resting Brain: Simultaneous Representation of Two Alpha Related Processes in Humans. PLoS ONE, 3(12), e3984. DOI:10.1371/journal.pone.0003984

Bergström, Z. M., Vogelsang, D. A., Benoit, R. G., & Simons, J. S. (2015). Reflections of Oneself: Neurocognitive Evidence for Dissociable Forms of Self-Referential Recollection. Cerebral Cortex (New York, N.Y.□: 1991), 25(9), 2648–57. DOI:10.1093/cercor/bhu063

Botvinick, M. 2004. Probing the neural basis of body ownership. Science, 305, 782–783. doi: 10.1126/science.1101836

Botvinick, M., & Cohen, J. (1998). Rubber hands “feel” touch that eyes see. Nature, 391(6669), 756–756. DOI:10.1038/35784

Braun, N., Debener, S., Spychala, N., Bongartz, E., Sörös, P., Müller, Helge H. O., Philipsen, A. 2018. The senses of agency and ownership: A Review. Frontiers in Psychology 9:535. doi: 10.3389/fpsyg.2018.00535

Brookes, M. J., Woolrich, M., Luckhoo, H., Price, D., Hale, J. R., Stephenson, M. C.,… Morris, P. G. (2011). Investigating the electrophysiological basis of resting state networks using magnetoencephalography. Proceedings of the National Academy of Sciences, 108(40), 16783–16788. DOI:10.1073/pnas.1112685108

Buzsáki, G., & Draguhn, A. (2004). Neuronal oscillations in cortical networks. Science (New York, N.Y.), 304(5679), 1926–9. DOI:10.1126/science.1099745

Buzsáki, G. 2019. The Brain from Inside Out. New York: Oxford University Press.

Canolty, R. T., Edwards, E., Dalal, S. S., Soltani, M., Nagarajan, S. S., Kirsch, H. E.,… Knight, R. T. (2006). High Gamma Power Is Phase-Locked to Theta Oscillations in Human Neocortex. Science, 313(5793), 1626–1628. DOI:10.1126/science.1128115

Cecere, R., Rees, G., & Romei, V. (2015). Individual differences in alpha frequency drive crossmodal illusory perception. Current Biology□: CB, 25(2), 231–5. DOI:10.1016/j.cub.2014.11.034

Clayton, M. S., Yeung, N., Cohen K. R. 2018. The many characters of visual alpha oscillations. European Journal of Neuroscience Oct;48(7):2498–2508. doi: 10.1111/ejn.13747.

Costantini, M., Robinson, J., Migliorati, D., Donno, B., Ferri, F., & Northoff, G. (2016). Temporal limits on rubber hand illusion reflect individuals? temporal resolution in multisensory perception. Cognition, 157, 39–48. DOI:10.1016/j.cognition.2016.08.010

Costantini, M., Robinson, J., Migliorati, D., Donno, B., Ferri, F., & Northoff, G. (2016). Temporal limits on rubber hand illusion reflect individuals’ temporal resolution in multisensory perception. Cognition, 157, 39–48. DOI:10.1016/j.cognition.2016.08.010

Cohen, M.X. (2014). Analyzing neural time series data: theory and practice. Massachusetts Institute of Technology.

Critchley, H. D., Botan, V., Ward, J. 2021. Absence of reliable physiological signature of illusory body ownership revealed by fine grained autonomic measurement during the rubber hand illusion. PLoS ONE 16(4): e0237282. https://doi.org/10.1371/journal.pone.0237282

de Haan, A. M., Van Stralen, H. E., Smit, M., Keizer, A., Van der Stigchel, S., Dijkerman, H. C., 2017. No consistent cooling of the real hand in the rubber hand illusion. Acta Psychologica 179, 68–77.

della Gatta, F., Garbarini, F., Puglisi, G., Leonetti, A., Berti, A., Borroni, P. 2016. Decreased motor cortex excitability mirrors own hand disembodiment during the rubber hand illusion. eLife 2016;5:e14972 DOI:10.7554/eLive.14972

de Vignemont, F. 2018. Mind the body: An exploration of bodily self-awareness. Oxford: Oxford University Press.

Delorme, A., & Makeig, S. (2004). EEGLAB: an open source toolbox for analysis of single-trial EEG dynamics including independent component analysis. Journal of Neuroscience Methods, 134(1), 9–21. DOI:10.1016/j.jneumeth.2003.10.009

Dienes, Z., Palfi, B., & Lush, P. 2020. Controlling phenomenology by being unaware of intentions. https://doi.org/10.31234/osf.io/4zw6g

Durgin, F. H., Evans, L., Dunphy, N., Klostermann, S. & Simmons, K. (2007) Rubber hands feel the touch of light. Psychological Science, 18, 152–157. doi:10.1111/j.1467-9280.2007.01865.x

Ehrsson, H. H. (2012). The concept of body ownership and its relation to multisensory integration. In B. E. Stein (Ed.) The new handbook on multisensory processing. Cambridge, MA: The MIT Press. pp. 775–793.

Ehrsson HH, Spence C, Passingham RE. 2004. That’s my hand! Activity in pre-motor cortex reflects feeling of ownership of a limb. Science 5, 303–312. doi: 10.1126/science.1097011.

Fingelkurts, A. A., Fingelkurts, A. A., Bagnato, S., Boccagni, C., & Galardi, G. (2012). EEG oscillatory states as neuro-phenomenology of consciousness as revealed from patients in vegetative and minimally conscious states. Consciousness and Cognition, 21(1), 149–169. doi:10.1016/j.concog.2011.10.004

Freeman, W. J., & Rogers, L. J. (2002). Fine temporal resolution of analytic phase reveals episodic synchronization by state transitions in gamma EEGs. Journal of Neurophysiology, 87(2), 937–45. DOI: 10.1152/jn.00254.2001

Germine, L., Benson, T. L., Cohen, F., & Hooker, C. I. (2013). Psychosis-proneness and the rubber hand illusion of body ownership. Psychiatry Research, 207(1-2), 45–52. DOI:10.1016/j.psychres.2012.11.022

Groppe, D. M., Urbach, T. P., & Kutas, M. (2011). Mass univariate analysis of event-related brain potentials/fields I: A critical tutorial review. Psychophysiology, 48(12), 1711–1725. DOI: 10.1111/j.1469-8986.2011.01273.x

Gusnard, D. A., Adbudak, E., Shulman, G. L., Raichle, M. E. 2001. Medial prefrontal cortex and self-referential mental activity: Relation to a default mode of brain function. Proceedings of the National Academy of Sciences 98, 7, 4259–4264.

Hanslmayr, S., Aslan, A., Staudigl, T., Klimesch, W., Herrmann, C. S., & B?uml, K.-H. (2007). Prestimulus oscillations predict visual perception performance between and within subjects. NeuroImage, 37(4), 1465–1473. DOI:10.1016/j.neuroimage.2007.07.011

Howells, F. M., Temmingh, H. S., Hsieh, J. H. van Dijen, A. V., Baldwin, D., Stein, D. J. 2018. Electroencephalographic delta/alpha frequency activity differentiates psychotic disorders: a study of schizophrenia, bipolar disorder and methamphetamine-induced psychotic disorder. Translational Psychiatry 8:75. DOI 10.1038/s41398-018-105-y.

Hutchinson, B. T., Pammer, K., Jack, B. 2021. Pre-stimulus alpha predicts inattentional blindness. Consciousness and Cognition Jan;87:103034. DOI: 10.1016/j.concog.2020.103034.

Jensen MP, Adachi T, Hakimian S. 2016. Brain Oscillations, Hypnosis, and Hypnotizability. American Journal of Clinical Hypnosis. 57, 3, 230–253. doi: 10.1080/00029157.2014.976786.

Kammers, M. P. M., Is this hand for real? Attenuation of the rubber hand illusion by transcranial magnetic stimulation over the inferior parietal lobule. J. Cogn. Neurosci. 21, 1311–1320 (2009).

Kanayama, N., Sato, A., & Ohira, H. 2009. The role of gamma band oscillations and synchrony on rubber hand illusion and crossmodal integration. Brain and Cognition, 69, 19–29. doi: 10.1016/j.bandc.2008.05.001.

Karamacoska, D., Barry, R. J., & Steiner, G. Z. (2017). Resting state intrinsic EEG impacts on go stimulus-response processes. Psychophysiology, 54(6), 894–903. DOI:10.1111/psyp.12851

Kihlstrom, J. F. 2016. Hypnosis. In H. S. Friedman (Ed.) Encyclopedia of Mental Health 2^nd^ Edition. pp. 361–365.

Klimesch, W., Sauseng, P., & Hanslmayr, S. (2007). EEG alpha oscillations: The inhibition?timing hypothesis. Brain Research Reviews, 53(1), 63–88. DOI:10.1016/j.brainresrev.2006.06.003

Knyazev, G. G. (2013). EEG Correlates of Self-Referential Processing. Frontiers in Human Neuroscience, 7. DOI:10.3389/fnhum.2013.00264

Knyazev, G. G. (2013). EEG Correlates of Self-Referential Processing. Frontiers in Human Neuroscience, 7, 264. DOI:10.3389/fnhum.2013.00264

Knyazev, G. G. (2013). Extraversion and anterior vs. posterior DMN activity during self-referential thoughts. Frontiers in Human Neuroscience, 6, 348. DOI:10.3389/fnhum.2012.00348

Knyazev, G. G., Savostyanov, A. N., Volf, N.V., Liou, M., & Bocharov, A.V. (2012). EEG correlates of spontaneous self-referential thoughts: A cross-cultural study. International Journal of Psychophysiology, 86(2), 173–181. DOI:10.1016/j.ijpsycho.2012.09.002

Kraus B, Salvador CE, Kamikubo A, et al. 2021. Oscillatory alpha power at rest reveals an independent self: A cross-cultural investigation. Biological Psychology. May;163:108118. DOI: 10.1016/j.biopsycho.2021.108118.

Lane, T. J. 2014 When actions feel alien: An explanatory model. In Hung, T. W. Ed. Communicative Action. Springer Science+Business Media. pp. 53–74.

Lane, T. J. 2015 Self, belonging, and conscious experience: A critique of subjectivity theories of consciousness. In R. Gennaro, Ed. Disturbed Consciousness: New Essays on Psychopathology and Theories of Consciousness. Cambridge, MA: MIT Press. pp. 103–140.

Lane, T. J. 2020. The minimal self hypothesis. Consciousness and Cognition. 2020 Oct;85:103029. doi: 10.1016/j.concog.2020.103029.

Lane T, Duncan NW, Cheng T, Northoff G. 2016. The Trajectory of Self. Trends in Cognitive Science. 20, 7, 481–482. doi: 10.1016/j.tics.2016.03.004.

Lane, T., Yeh, S.-L., Tseng, P., & Chang, A.-Y. 2017. Timing disownership experiences in the rubber hand illusion. Cognitive Research: Principles and Implications, 2(1), 4. DOI:10.1186/s41235-016-0041-4

Lehmann, D., Faber, P. L., Achermann, P., Jeanmonod, D., Gianotti, L. R., & Pizzagalli, D. (2001). Brain sources of EEG gamma frequency during volitionally meditation-induced, altered states of consciousness, and experience of the self. Psychiatry Research, 108(2), 111–21. DOI: 10.1016/S0925-4927(01)00116-0

Lenggenhager, B., Halje, P., & Blanke, O. (2011). Alpha band oscillations correlate with illusory self-location induced by virtual reality. European Journal of Neuroscience, 33(10), 1935–1943. DOI:10.1111/j.1460-9568.2011.07647.x

Limanowski, J., Lutti, A., Blankenburg, F. 2014. The extrastriate body area is involved in illusory limb ownership. Neuroimage 86, 514–524.

Limanowski, J., & Blankenburg, F. (2016). Integration of Visual and Proprioceptive Limb Position Information in Human Posterior Parietal, Premotor, and Extrastriate Cortex. Journal of Neuroscience, 36(9), 2582–2589. DOI:10.1523/JNEUROSCI.3987-15.2016

Lira, M., Pantaleão, F. N., de Souza Ramos, C. G., Boggio, P. S. 2018. Anodal transcranial direct current stimulation over the posterior parietal cortex reduces the onset time to the rubber hand illusion and increases the body ownership. Experimental Brain Research 236, 11, 2935–2943. doi: 10.1007/s00221-018-5353-9.

Lloyd, D. M. (2007). Spatial limits on referred touch to an alien limb may reflect boundaries of visuo-tactile peripersonal space surrounding the hand. Brain and Cognition, 64, 104–109.

Longo, M. R., Schüür, F., Kammers, M. P. M., Tsakiris, M., & Haggard, P. (2008). What is embodiment? A psychometric approach. Cognition, 107(3), 978–98. DOI:10.1016/j.cognition.2007.12.004

Lu, H., Zuo, Y., Gu, H., Waltz, J. A., Zhan, W., Scholl, C. A.,… Stein, E. A. (2007). Synchronized delta oscillations correlate with the resting-state functional MRI signal. Proceedings of the National Academy of Sciences, 104(46), 18265–18269. DOI:10.1073/pnas.0705791104

Lush, P., Moga, G., McLatchie, N., Dienes, Z. 2018. The Sussex-Waterloo Scale of Hypnotizability (SWASH): measuring capacity for altering conscious experience, Neuroscience of Consciousness, Volume 2018, Issue 1, 2018, niy006, https://doi.org/10.1093/nc/niy006

Lush, P., Botan, R. B., Scott, R. B., Seth, A. K., Ward, J. Dienes, Z. 2020. Trait phenomenological control predicts experience of mirror synaesthesia and the rubber hand illusion. Nature Communications 11,1, 4853. doi:10.1038/s41467-020-18591-6

Lush, P., Seth, A., & Dienes, Z. (2021, May 7). Demand characteristics confound asynchronous control conditions in indirect measures of the rubber hand illusion. https://doi.org/10.31234/osf.io/w67xc

Mantini, D., Perrucci, M. G., DelGratta, C., Romani, G. L., & Corbetta, M. (2007). Electrophysiological signatures of resting state networks in the human brain. Proceedings of the National Academy of Sciences, 104(32), 13170–13175. DOI:10.1073/pnas.0700668104

Marotta, A., Tinazzi, M., Cavedini, C., Zampini, M., Fiorio, M. 2016. Individual Differences in the Rubber Hand Illusion Are Related to Sensory Suggestibility. PLoS ONE 11,12: e0168489. https://doi.org/10.1371/journal.pone.0168489

Newson, J. J, Thiagarajan, T.C. 2019. EEG Frequency Bands in Psychiatric Disorders: A Review of Resting State Studies. Frontiers in Human Neuroscience 2019; 12:521. doi:10.3389/fnhum.2018.00521.

Northoff, G., & Bermpohl, F. (2004). Cortical midline structures and the self. Trends in Cognitive Sciences, 8(3), 102–107. doi:10.1016/j.tics.2004.01.004

Ocklenburg, S., Ruther, N., Peterburs, J., Pinnow, M., & Gunturkun, O. (2011). Laterality in the rubber hand illusion. Laterality: Asymmetries of Body, Brain and Cognition, 16(2), 174–187. DOI:10.1080/13576500903483515

Park, H. D., Blanke, O. 2019. Coupling inner and outer body for self-consciousness. Trends in cognitive sciences 23, 5, 377–388.

Peled, A., Ritsner, M., Hirschmann, S., Geva, A. B., & Modai, I. (2000). Touch feel illusion in schizophrenic patients. Biological Psychiatry, 48(11), 1105–8. DOI: 10.1016/S0006-3223(00)00947-1

Peled, A., Pressman, A., Geva, A. B., & Modai, I. (2003). Somatosensory evoked potentials during a rubber-hand illusion in schizophrenia. Schizophrenia Research 64, 2-3, 157–63. DOI: 10.1016/S0920-9964(03)00057-4

Petkova, V., Bj?rnsdotter, M., Gentile, G., Jonsson, T., Li, T.-Q., & Ehrsson, H. □?Henri. (2011). From Part- to Whole-Body Ownership in the Multisensory Brain. Current Biology, 21(13), 1118–1122. DOI:10.1016/j.cub.2011.05.022

Peviani, V., Magnani, F. G., Ciricugno, A., Vecchi, T., Bottini, G. 2018. Rubber hand illusion survives Ventral premotor area inhibition: A rTMS study. Neuropsychologia 120, 18–24.

Piantoni, G., Romeijn, N., Gomez-Herrero, G., Van Der Werf, Y. D., Van Someren, E. J. W. Alpha power predicts persistence of bistable perception. Scientific Reports 7:5208. DOI:10.1038/s41598-017-05610-8.

Powell, J. L., Lewis, P. A., Dunbar, R. I. M., Garc?a-Fi?ana, M., & Roberts, N. (2010). Orbital prefrontal cortex volume correlates with social cognitive competence. Neuropsychologia, 48(12), 3554–3562. DOI:10.1016/j.neuropsychologia.2010.08.004

Qin, P., Grimm, S., Duncan, N. W., Fan, Y., Huang, Z., Lane, T. et al. 2016. Spontaneous activity in default-mode network predicts ascription of self-relatedness stimuli. Social cognitive and affective neuroscience 11, 4, 693–702.

Ranlund, S., Nottage, J., Shaikh, M., Dutt, A., Constante, M., Walshe, M.,… Bramon, E. (2014). Resting EEG in psychosis and at-risk populations ? A possible endophenotype? Schizophrenia Research, 153(1-3), 96–102. DOI:10.1016/j.schres.2013.12.017

Rao, I. S., Kayser, C. 2017. Neurophysiological correlates of the rubber hand illusion in Late Evoked and Alpha/Beta Band Activity. Frontiers in Human Neuroscience 11: 377. doi:10.3389/fnhum.2017.00377

Riemer, M., Trojan, J., Beauchamp, M., Fuchs, X. 2019. The rubber hand universe: On the impact of methodological differences in the rubber hand illusion. Neuroscience & Biobehavioral Reviews 104, 268–280.

Roach, B. J., & Mathalon, D. H. (2008). Event-Related EEG Time-Frequency Analysis: An Overview of Measures and An Analysis of Early Gamma Band Phase Locking in Schizophrenia. Schizophrenia Bulletin, 34(5), 907–926. DOI:10.1093/schbul/sbn093

Romano, D., Maravita, A. & Perugini, M. (2021). Psychometric properties of the embodiment scale for the rubber hand illusion and its relation with individual differences. Scientific Reports 11, 5029. https://doi.org/10.1038/s41598-021-84595-x

Romei, V., Brodbeck, V., Michel, C., Amedi, A., Pascual-Leone, A., & Thut, G. 2008. Spontaneous fluctuations in posterior alpha-band EEG activity reflect variability in excitability of human visual areas. Cerebral Cortex (New York, N.Y. □: 1991), 18(9), 2010–8. DOI:10.1093/cercor/bhm229

Romei, V., Rihs, T., Brodbeck, V., & Thut, G. 2008. Resting electroencephalogram alpha-power over posterior sites indexes baseline visual cortex excitability. Neuroreport, 19(2), 203–8. DOI:10.1097/WNR.0b013e3282f454c4

Roy, M., Shohamy, D., & Wager, T. D. (2012). Ventromedial prefrontal-subcortical systems and the generation of affective meaning. Trends in Cognitive Sciences, 16(3), 147–56. DOI:10.1016/j.tics.2012.01.005

Roy, M., Shohamy, D., & Wager, T. D. (2012). Ventromedial prefrontal-subcortical systems and the generation of affective meaning. Trends in Cognitive Sciences, 16(3), 147–156. DOI:10.1016/j.tics.2012.01.005

Samaha, J., & Postle, B. R. (2015). The Speed of Alpha-Band Oscillations Predicts the Temporal Resolution of Visual Perception. Current Biology□: CB, 25(22), 2985–90. DOI:10.1016/j.cub.2015.10.007

Smallwood, J., Schooler, J. W. 2015. The science of mind wandering: Empirically navigating the stream of consciousness. Annual Review of Psychology 66, 487–518.

Stokes, M., Buschman, T.J., and Miller, E.K. (2017). Dynamic coding for flexible cognitive control. The Wiley Handbook of Cognitive Control, Edited by Tobias Egner, John Wiley & Sons, (Chichester, West Sussex, UK).

Swinkels, L. M. J., Veling, H., Dijksterjuis, A., van Schie, H. T. 2020. Availability of synchronous information in an additional sensory modality does not enhance the full body illusion. Psychological Research 2020 Jul 27. doi: 10.1007/s00426-020-01396-z.

Tastevin, J. (1937). En parent de l’experience d’Aristotle. L’Encephale, I, 57–84.

Thakkar, K. N., Nichols, H. S., McIntosh, L. G., & Park, S. (2011). Disturbances in body ownership in schizophrenia: evidence from the rubber hand illusion and case study of a spontaneous out-of-body experience. PloS One, 6(10), e27089. DOI:10.1371/journal.pone.0027089

Thut, G., Nietzel, A., Brandt, S. A., & Pascual-Leone, A. (2006). ?-Band Electroencephalographic Activity over Occipital Cortex Indexes Visuospatial Attention Bias and Predicts Visual Target Detection. Journal of Neuroscience, 26(37), 9494–9502. DOI:10.1523/JNEUROSCI.0875-06.2006

Tsakiris, M. 2010. My body in the brain: A neurocognitive model of body-ownership. Neuropsychologia, 48(3), 703–712. DOI:10.1016/j.neuropsychologia.2009.09.034

Tsakiris, M. 2017. The multi-sensory basis of the self: From body to identity to others. The quarterly journal of experimental psychology 70, 4, 597–609.

Tsakiris, M., & Haggard, P. (2005). The Rubber Hand Illusion Revisited: Visuotactile Integration and Self-Attribution. Journal of Experimental Psychology: Human Perception and Performance, 31(1), 80–91. DOI:10.1037/0096-1523.31.1.80

Tsakiris, M., Hesse, M. D., Boy, C., Haggard, P., & Fink, G. R. (2007). Neural signatures of body ownership: a sensory network for bodily self-consciousness. Cerebral Cortex (New York, N.Y.□: 1991), 17(10), 2235–44. DOI:10.1093/cercor/bhl131

Vanhatalo, S., Palva, J. M., Holmes, M. D., Miller, J. W., Voipio, J., & Kaila, K. (2004). Infraslow oscillations modulate excitability and interictal epileptic activity in the human cortex during sleep. Proceedings of the National Academy of Sciences, 101(14), 5053–5057. DOI:10.1073/pnas.0305375101

Wolff, M. J., Jochim, J., Akyürek, E. G., & Stokes, M. G. (2017). Dynamic hidden states underlying working-memory-guided behavior. Nature Neuroscience, 20(6), 864–871. DOI:10.1038/nn.4546

Yeh, S-L., Lane, T. J., Chang, A-Y., Chien, S-E. 2017. Switching to the rubber hand. Frontiers in Psychology: Cognitive Science DOI: 10.3389/fpsyg.2017.02172

Zeller, D., Litvak, V., Friston, K. J., & Classen, J. 2015. Sensory processing and the rubber hand illusion—an evoked potentials study. Journal of Cognitive Neuroscience, 27, 3, 573–582. DOI:10.1162/jocn_a_00705

